# Model for ring closure in ER tubular network dynamics

**DOI:** 10.1101/2021.11.18.469198

**Authors:** Ben Zucker, Gonen Golani, Michael M. Kozlov

**Author notes:** These authors equally contributed.

## Abstract

Tubular networks of endoplasmic reticulum (ER) are dynamic structures whose steady-state conformations are maintained by a dynamic balance between the persistent generation and vanishing of the network elements. While factors producing the ER tubules and inter-tubular junctions have been investigated, the mechanisms behind their elimination remained unknown. Here we addressed the ER ring closure, the process resulting in the tubule and junction removal through constriction of the network unit-cells into junctional knots followed by the knot remodeling into regular junctions. We considered the ring closure to be driven by the tension existing in ER membranes. We modeled, computationally, the structures of the junctional knots containing internal nanopores and analyzed their tension dependence. We predicted an effective interaction between the nanopores facilitating the knot tightening and collapse of additional network unit cells. We analyzed the process of the pore sealing through membrane fission resulting in formation of regular junctions. Considering the hemi-fission as the rate-limiting stage of the fission reaction, we evaluated the membrane tensions guarantying the spontaneous character of the pore sealing. We concluded that feasible membrane tensions explain all stages of the ER ring closure.

## Introduction

Endoplasmic reticulum (ER) is one of the most vital membrane bound organelles of eukaryotic cells serving as a platform for protein synthesis, folding, modification and transport to intra-cellular destination(Alberts et al., 2017, chap. 12). The ER functions determine its structure consisting of the double-membrane nuclear envelope, and the peripheral ER, which spans most of the cytoplasm of mammalian cells(Westrate et al., 2015). The peripheral ER includes 30-50nm thick flat membrane sheets and a highly intricate network of membrane tubules with 25-90 nm diameters interconnected by three-way junctions(Schroeder et al., 2018; Terasaki, 2018; Voeltz et al., 2002; Westrate et al., 2015). In spite of their complex architecture and topological features, the membranes of all ER elements are interconnected into a continuous system that encloses a common luminal space.

Several decade-long studies by the methods of diffraction-limited optical(Lee and Chen, 1988; Terasaki et al., 1986), electron(Palade, 1956), and, more recently, super-resolution(Nixon-Abell et al., 2016; Schroeder et al., 2018) microscopies have revealed a multiscale character of the peripheral ER architecture exhibiting a macroscopic and nanoscopic levels of organization(Westrate et al., 2015). The ER nanoscopic structures described so far include ER tubular matrices(Nixon-Abell et al., 2016), sheet nanoholes(Schroeder et al., 2018), and ER to Golgi transport intermediates(Weigel et al., 2021), all characterized by an intrinsic length scale in the range between few tens and 100 nm. The nanoscopic structures appear to represent mechanically equilibrium membrane conformations. The macroscopic ER structures include stacks of micron-large sheets interconnected by helicoidal junctions(Terasaki et al., 2013), and irregular polygonal networks of tubules with a typical unit-cell size of several microns that are in the focus of this study.

The macroscopic tubular networks, exhibit an, essentially, non-equilibrium dynamic behavior, which includes continuous emerging and disappearance of junctions and tubules on the background of their perpetual motion(Lee and Chen, 1988). The junctions and tubules are generated through the new tubule branching off and fusion with the tubules already existing within the network(Dabora and Sheetz, 1988; Lee and Chen, 1988). This leads to formation of new network unit-cells. Disappearance of the junctions and tubules is mediated by the process referred to as the ring closure, whose essence is a contraction followed by vanishing of the network unit-cells(Lee and Chen, 1988; Powers et al., 2017). The contraction stage results in transformation of a polygonal unit-cell, regardless of its initial size, into a small “knot”, which replaces one of the initial inter-tubular junctions. In the following, we will refer to this structure as the junctional knot. Normally, a junctional knot transforms into a regular junction with no noticeable swelling or discontinuity(Lee and Chen, 1988), thus, completing the ring closure. In addition to the ring closure, a direct scission of ER tubules has been reported(Espadas et al., 2019), but appears to happen too rarely to play a considerable role in the network dynamics(Wang et al., 2016). In spite of the persistent generation and disappearance of the network elements, the overall numbers of the junctions, tubules and unit-cells in the macroscopic tubular network do not change in time. The steady-state conformations of ER macroscopic networks are commonly attributed to the dynamic balance between the tubule branching-fusion and the ring closure events(Lee and Chen, 1988).

Understanding the physical mechanisms behind the dynamics of the macroscopic ER networks, requires knowledge of a minimal set of physical factors, which drive the competing processes of creation and disintegration of the network elements. Based on the i*n vitro* studies(Wang and Rapoport, 2019) and live cell observations, there are three such factors. The proteins of the reticulon/REEP families generate and stabilize ER tubules by producing the required membrane curvature(Hu et al., 2008). The Atlastin family proteins create the network junctions by mediating fusion of ER tubules(Powers et al., 2017). Pulling forces applied to the ER membranes by the cytoskeleton appear to be necessary, on one hand, for the tubular branching, mobility, and maintenance of the overall extended network conformations, and, on the other, for the ring closure and the related elimination of the network elements. This is supported by the observations that disassembly of the intracellular force-generating machinery based on microtubules and the associated molecular motors results in stopping all kinds of movement of the ER network elements including the unit-cell contraction and the related ring closure(Terasaki et al., 1986). Also the reconstituted ER tubular networks, which exhibited the major structural and dynamic features of the intra-cellular ER networks including the ring closure(Powers et al., 2017; Wang and Rapoport, 2019), were, most probably, subjected to pulling forces originating from such factors as the network adhesion to the external substrate and the thermal convection of the surrounding liquid(15).

While micromechanics of the protein-mediated membrane curving and fusion involved in generation of the ER network elements have been thoroughly investigated(Wang et al., 2021; Wang and Rapoport, 2019), the physical mechanisms behind the disintegration of the network tubules and junctions trough the ring closure have never been addressed and represent the subject of this work.

Here we analyze the formation of the junctional knots and their remodeling into regular junctions as the crucial stages of the ER ring closure. We propose that the major factor driving all steps of these processes is the membrane tension. We computationally determine the experimentally inaccessible configurations of the junctional knots containing internal pores and resulting from constriction of one or multiple network unit-cells. We analyze fission of the hourglass-like membrane necks forming the pores rims, which mediates the pore sealing and constitutes the essence of the knot remodeling into the regular junctions. We consider the fission to proceed through an intermediate stage of hemi-fission representing the rate-limiting stage of the process. We compute the energy of the hemi-fission structure determining the energy barrier of the fission reaction and evaluate the tensions necessary to eliminate this barrier.

## Model

We consider the process of ring closure illustrated in (Fig. 1A-H) as consisting of two stages: constriction of an ER network unit cell into a junctional knot (Fig. 1A-D), and transformation of the latter into a regular junction (Fig.1 E-H).

**Figure 1.**
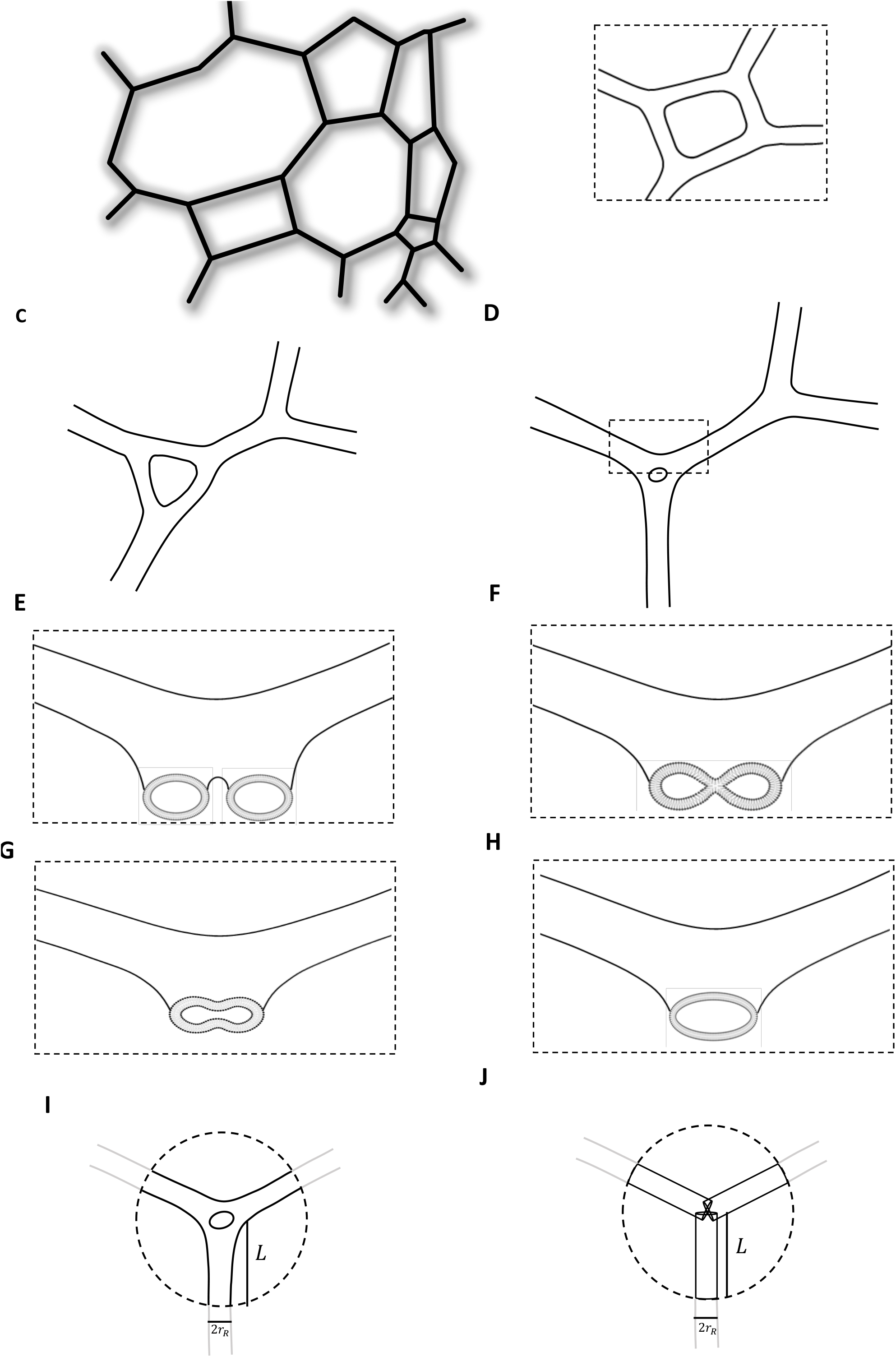
Model for the pathway of the ER ring closure. (A-D) Formation of a junctional knot as a result of the ring constriction. (A) Segment of macroscopic network of ER tubules. (B) A network unit-cell with four junctions. (C) Formation of a triangular unit cell as a result of dynamic evolution of the four-junction unit cell. (D) Constriction of the triangular unit cell into a junctional knot. (E-H) Transformation of a junctional knot into a regular junction trough the pore sealing mediated by membrane fission. (E) View of the junctional knot cross-section indicated in (H). (F) View of the cross-section through the hemi-fission stalk. (G) and (H) represents sequential stages of post fission relaxation. (I) The system under consideration in contact with the reservoir with the dashed circle indicating the reservoir boundary, *r_R_* is the radius of the reservoir tubules, *L* is the distance from the system center to the boundaries with the reservoir. (J) The hypothetical reference system.

We assume the unit-cell constriction (Fig.1 A-D) to be driven by the membrane tension. This is based on the known dynamic property of random networks of stressed lines interconnected by mobile three-way junctions(Stavans, 1993). A simple balance analysis of the tension-related forces acting on the network junctions shows that all polygonal unit-cells with less than six vertices (i.e. pentagonal, quadratic and triangular) are mechanically unstable and tend to collapse by constriction(Stavans, 1993). The computational simulations of such network behavior demonstrated the events of the unit cell constriction and other features of the ER network dynamics(Lin et al., 2017). Examples of such simulations are presented in (Supplementary Movie). As follows from these and other simulations, polygonal unit-cells transform into equilateral triangular unit cells, which then undergo collapse into junctional knots (Fig.1 A-D).

Here we focus on the second stage of the ring closure (Fig.1 E-H), the remodeling of the junctional knots into regular junctions. The analysis is based on comparison of the elastic energy of the junctional knot membrane with that of the structures formed at the intermediate and final steps of the remodeling process.

We consider the junctional knot energy and the energy of the final regular junction to consist of two contributions: the energy of membrane bending, *F_B_*, and the energy of membrane tension, *F_T_*,

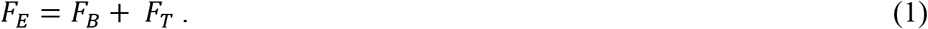

The bending energy, *F_B_*, is associated with the shape of the membrane midplane that is characterized at each point by the mean, *J*, and Gaussian, *K*, curvatures(Spivak, 1999). We use Helfrich model of membrane bending elasticity(Helfrich, 1973) according to which the membrane material parameters setting the value of the bending energy are the spontaneous curvature, *J_S_*, the bending modulus, *κ*, and the modulus of Gaussian curvature, 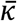 (Helfrich, 1973). The spontaneous curvature, *J_S_*, which accounts for the intrinsic tendency of the membrane to acquire a bent shape and determines the intra-membrane torque, is generated for ER tubular membranes by the curvature generating proteins of the reticulon/REEP family and has values in the 0.01 − 0.1nm^−1^ range, as inferred from the measurements of the ER tubule diameters(Schroeder et al., 2018; Terasaki, 2018; Voeltz et al., 2002). The bending modulus, *κ*, which sets the scale of the bending energy, while varying in a certain range(Dimova, 2014), has a typical value of, *κ* ≈ 10^−19^Joule(Dimova, 2014). The modulus of Gaussian curvature, 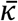, whose direct measurements have been, largely, prohibited by geometrical restrictions, is, typically, negative and has been estimated to have values in the range between 0 and −*κ* [see e.g.(Templer et al., 1998; Terzi et al., 2019)].

The tension energy, *F_T_*, is associated with the membrane area, *A*, and the system parameter setting the scale of this energy is the tension value, *γ*.

We define the system energy as the thermodynamic work of creating the junctional knot (or the regular junction) out of an effective membrane reservoir represented by the surrounding tubular ER. Based on the ER structure, we consider the reservoir to consist of cylindrical membrane tubules characterized by a spontaneous curvature, *J_S_*, and subject to a tension, *γ*. The radius, *r_R_*, and the corresponding mean curvature, *J_R_*, of the reservoir tubules are given by

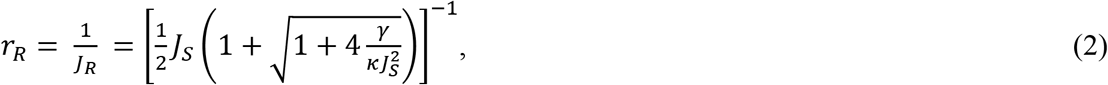

as set by the mechanical equilibrium between the tension and the intra-membrane torque (see SI A)). We consider the junctional knot to be connected to the reservoir by three tubular arms, which merge with the reservoir along circular border-lines of radius, *r_R_* (Eq. 2), and denote the distance from the border-lines to the unit-cell center by *L* (Fig. 1I). The boundary conditions imposed on membrane shape at the border-lines require a smooth transition between the membranes of the tubular arms and those of the reservoir. Taking the distance, *L*, to be substantially larger than the characteristic length-scale of the system, 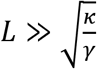 (see SI A)), we guarantee a negligibly small influence of the boundary conditions on the junctional knot configurations.

The bending energy of shaping a unit area of the reservoir membrane into a unit area element of the junctional knot (or the regular junction) is given by

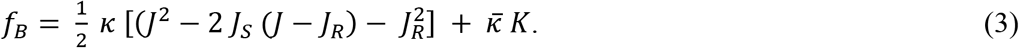

The total bending energy, *F_B_*, is obtained by integration of *f_B_* over the system area *A_T_*, which includes the area of the junctional knot (or the regular junction) and that of the connecting arms,

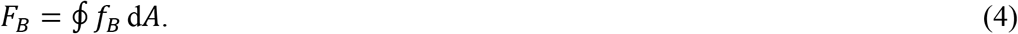

The tension energy is given by,

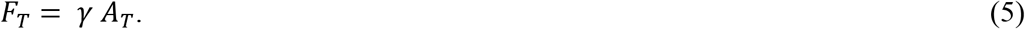

It is convenient to relate the energy of the junctional knot (or the regular junction), *F_E_*, to that of a hypothetical reference state, *F*_0_, which we define as consisting of three homogeneous cylindrical membrane tubules of radius *r_R_* (Eq. 2) pulled out of the reservoir through the boundary lines up to the center of the unit cell, i.e., each having the length *L* (Fig. 1J). The relative energy, *F_R_* = *F_E_* – *F*_0_, does not depend on the arbitrarily chosen length, *L*, provided that the latter is sufficiently large, and characterizes solely the conformation of the knot membrane *per se*.

By considering the energy of the intermediate stage of the junctional knot remodeling emerging as a result of hemi-fission of the pore rim, we account, in addition to the elastic energy above, for the energy of tilting and splaying the hydrocarbon chains of lipid molecules forming the membrane of the core of the hemi-fission structure. The description of the model used for evaluation of the tilt and splay energy and the related elastic moduli of the membrane is presented in (SI B).

## Results

### Configurations and energies of junctional knots and regular junction

The configurations and energies of the junctional knots (and the regular junctions) were obtained by numerical minimization of the relative energy, *F_R_*, using the Brakke’s Surface Evolver program(Brakke, 1992). The examples of the resulting configurations computed for a specific tension value are presented in (Fig. 2A), and (Fig.2B) for a junctional knot and a regular junction, respectively. Whereas the junctional knot and the regular junction have similar dimensions and shapes of their external contours (Fig. 2A,B), the former differs from the latter by an hour-glass pore in the middle (Fig. 2A). This pore is a remnant of the micron large space confined within the initial network unit-cell before its contraction. The equilibrium size of the pore is set by the interplay of two counter-acting factors, the tension and the bending energy. The bending energy of the membrane forming the pore rim increases in the course of the pore constriction and, hence, resists this process. By contrast, the tension favors the pore tightening. Therefore, the radius of the pore waist decreases from 8nm to 4nm with increasing membrane tension from 0.1 mN/m to 0.5 mN/m (Fig.2C). For even larger tensions the pore radius tends to decrease beneath 4 nm which is the lower boundary of our calculation validity set by the membrane thickness.

**Figure 2.**
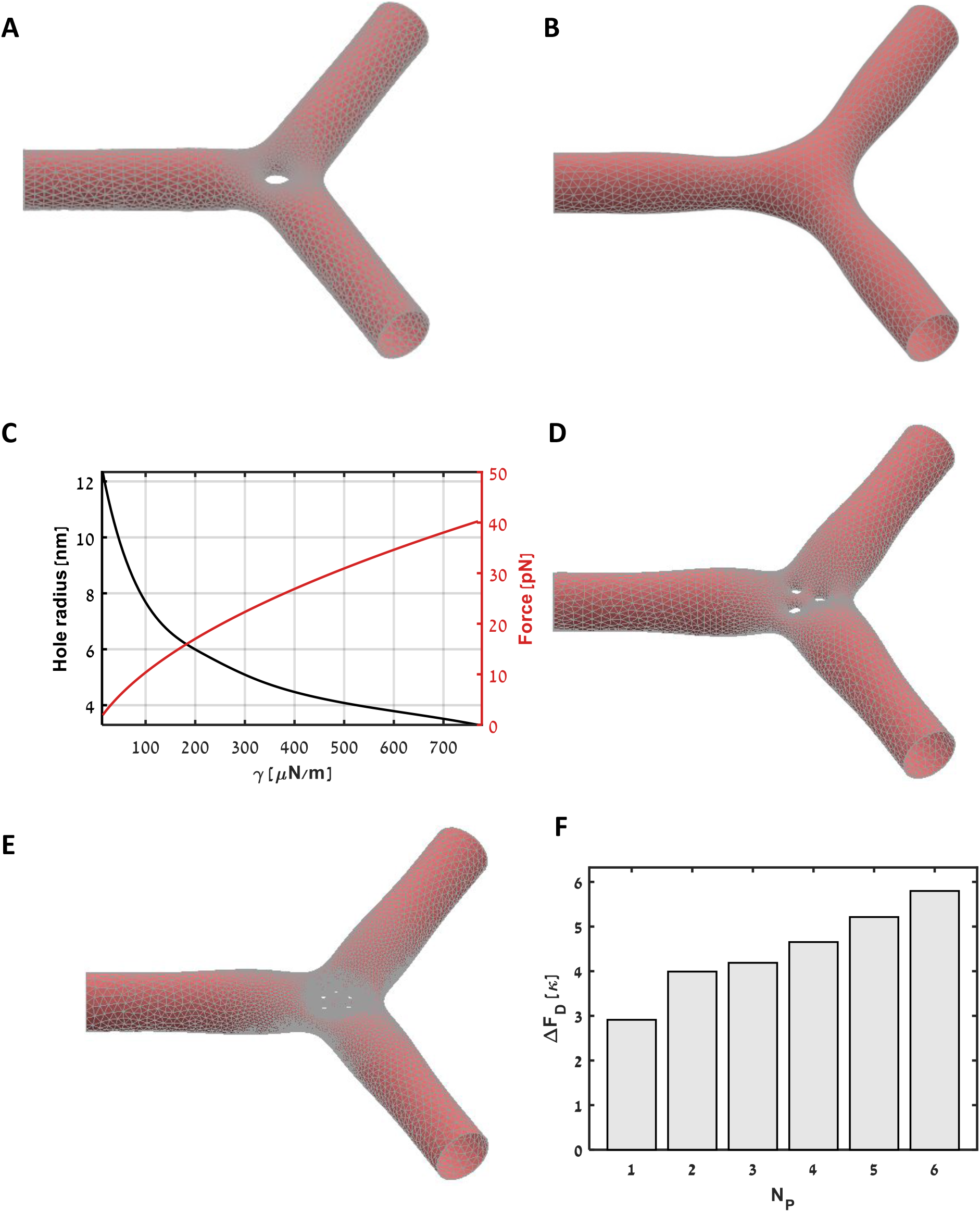
Computed configurations of junctional knots with different number of pores. (A) One pore. (B) No pore, which represents a regular junction. (C) The tension dependence of the pore radius in one-pore junctional knot. (D) Three pore junctional know. (E) Five pore junctional knot. (F) The energy of transition of one pore from the junctional knot to a tubule as a function of the initial number of pores in the junction. The used parameters: *κ* = 0.8 · 10^−19^Joule; 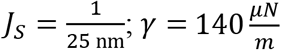 (except for (C)); *r_R_* = 15nm (except for (C)); *L* = 8.5 · *r_R_*.

We further analyzed the structures of junctional knots resulting from a sequential contraction of several triangular network unit-cells into the same junctional knot. To this end, we computed the optimal configurations of the junctional knots with various number of pores as presented in (Figs. 2D,E) for knots containing, respectively, three and five pores. As shown in (Fig.2 D,E), the pores are tightly packed in the center of the structure. The larger the pore number within a junctional knot, the smaller the pore radii under the same tension. This can be regarded as an effective tension-mediated attractive interaction between the pores within a junctional knot, which enables the pore contraction and tighter packing, and must facilitate embedding of additional pores.

Finally, we investigated the stability of the pore location within the junctional knot. For this purpose, we computed the change of the system energy resulting from displacement of one of the pores from the junctional knot into one of the tubules emerging from the junctional knot (Fig. 2F). As presented in (Fig. 2F), this event consumes energy irrespectively of the number of the pores within the junction meaning that the dense packing of the pores within a junctional knot is a stable configuration.

### Transformation of junctional knots into regular junctions through sealing of intra-knot pores

Here we analyze sealing of the intra-knot pores, which converts the junctional knots into regular junctions. This is a final step of the ring closure event, which reduces the overall number of junctions in the ER network. For a junctional knot containing one pore, the sealing completes the removal of two junctions from the network (Fig. 1E-H).

We consider the membrane fission mediating the sealing (Kozlovsky and Kozlov, 2003) to occur either within the hourglass-like membrane neck forming the rim of the pore, or within one of the short neck-like membrane elements forming the sides of the junctional knot and surrounding the pore. We propose that the factor driving the membrane fission is the membrane stress that accumulates within the junctional knot membrane under the tension-driven constriction of the pore and relaxes as a result of the pore sealing.

#### Favorability of the pore sealing

We start with analyzing the conditions required for the pore sealing to be energetically favorable, i.e., leading to decrease of the system energy. To this end we compare the energy, *F*_*E*0_, of a three-way tubular junction with no pores (Fig. 2B) with the energy, *F*_*E*1_, of a junctional knot with one pore (Fig. 2A).

The computational results for *F*_*E*0_, *F*_*E*1_ and for the energy released by the pore sealing, Δ*F_S_* = *F*_*E*1_ – *F*_*E*0_, are presented in (Fig. 3A,B) in dependence on the membrane tension, *γ*. According to (Fig. 3B), for non-vanishing tensions, the released energy, Δ*F_S_*, is positive and, hence, the sealing event is favorable within the considered parameter ranges. The energy release is a consequence of relaxation, as a result of the pore sealing, of the elastic stresses accumulated initially, within the junctional knot upon its tension-driven constriction. The tension reinforces the energetic favorability of the pore sealing since the amount of the released energy, Δ*F_S_*, increases with *γ* (Fig. 3B).

**Figure 3.**
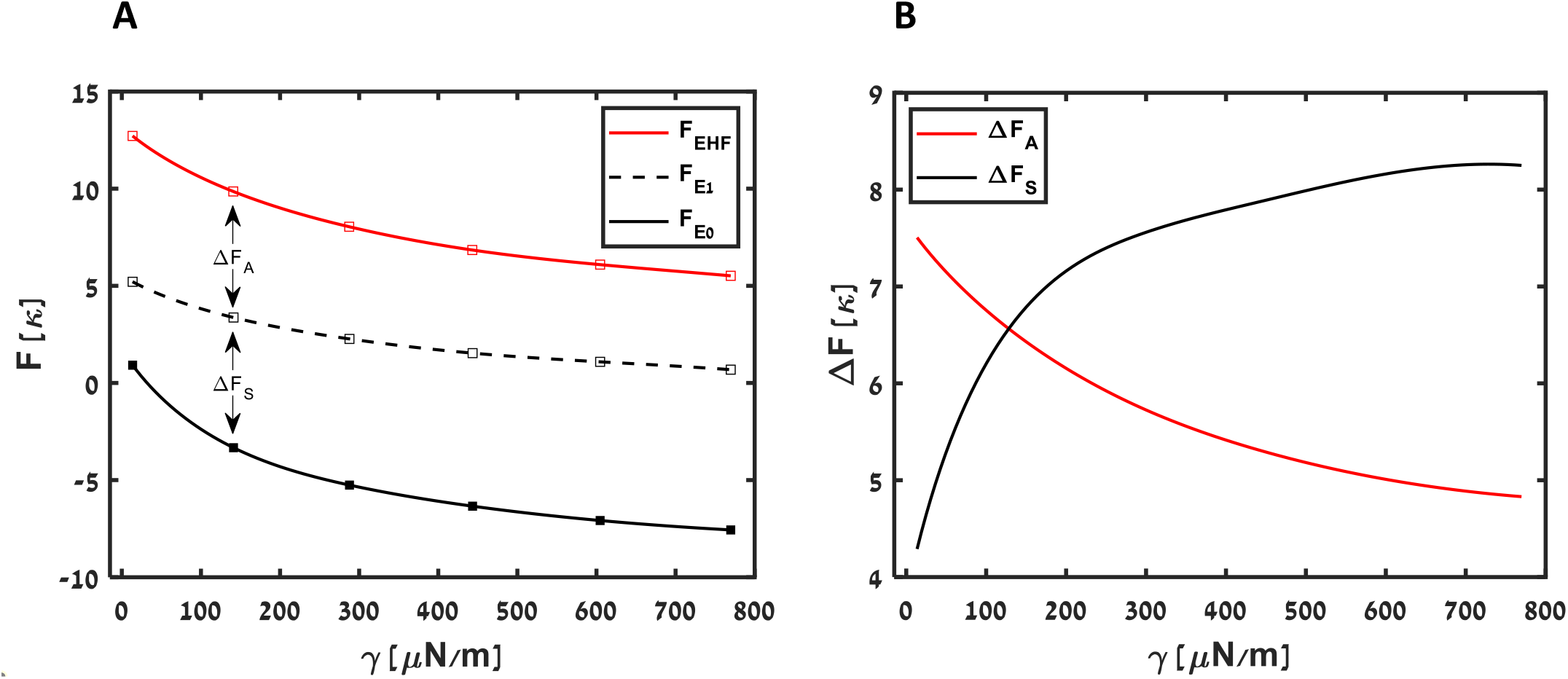
Energies of the intermediate stages of the ring closure as functions of tension. (A) The energies of the junctional knot with one pore (black line), the hemi-fission stage of the pore sealing (red line), the regular junction (dashed line). (B) The energy released as a result of the pore sealing, Δ*F_S_*, (black line), and the fission activation energy, Δ*F_A_*, (red line). *κ* = 0.8 · 10^−19^Joule; 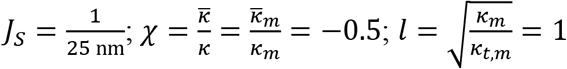.

#### Dependence of the rate-limiting stage of the pore sealing on the membrane tension

Finally, we address a possible kinetic hindrance of the pore sealing by an energy barrier represented by an intermediate state of the reaction. To this end, we consider a specific pathway of the membrane structural transformations leading from a junctional knot with one pore (Fig. 2A) to a regular junction (Fig. 2B).

First, we find the site within the junctional knot in which membrane fission occurs with a highest probability. We require the fission site to be a region encircling the membrane shape and characterized by a relatively high surface density of the elastic energy, *f_E_*, which includes the contributions from the bending energy, *f_B_* (Eq.2), and the reservoir tension, *γ*, such that

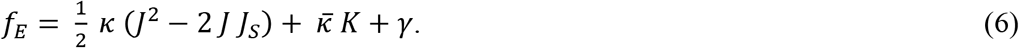

The computed distribution of *f_E_* along the surface of a junctional knot with one pore is presented in (Fig. 4A) for two limiting sets of the elastic parameters. In both cases, a continuous ring-like region of high or elevated energy density forms around the waist of the pore rim rather than the membrane necks surrounding the pore. Thus, we suggest that fission occurs in the middle of the membrane neck forming the pore rim.

**Figure 4.**
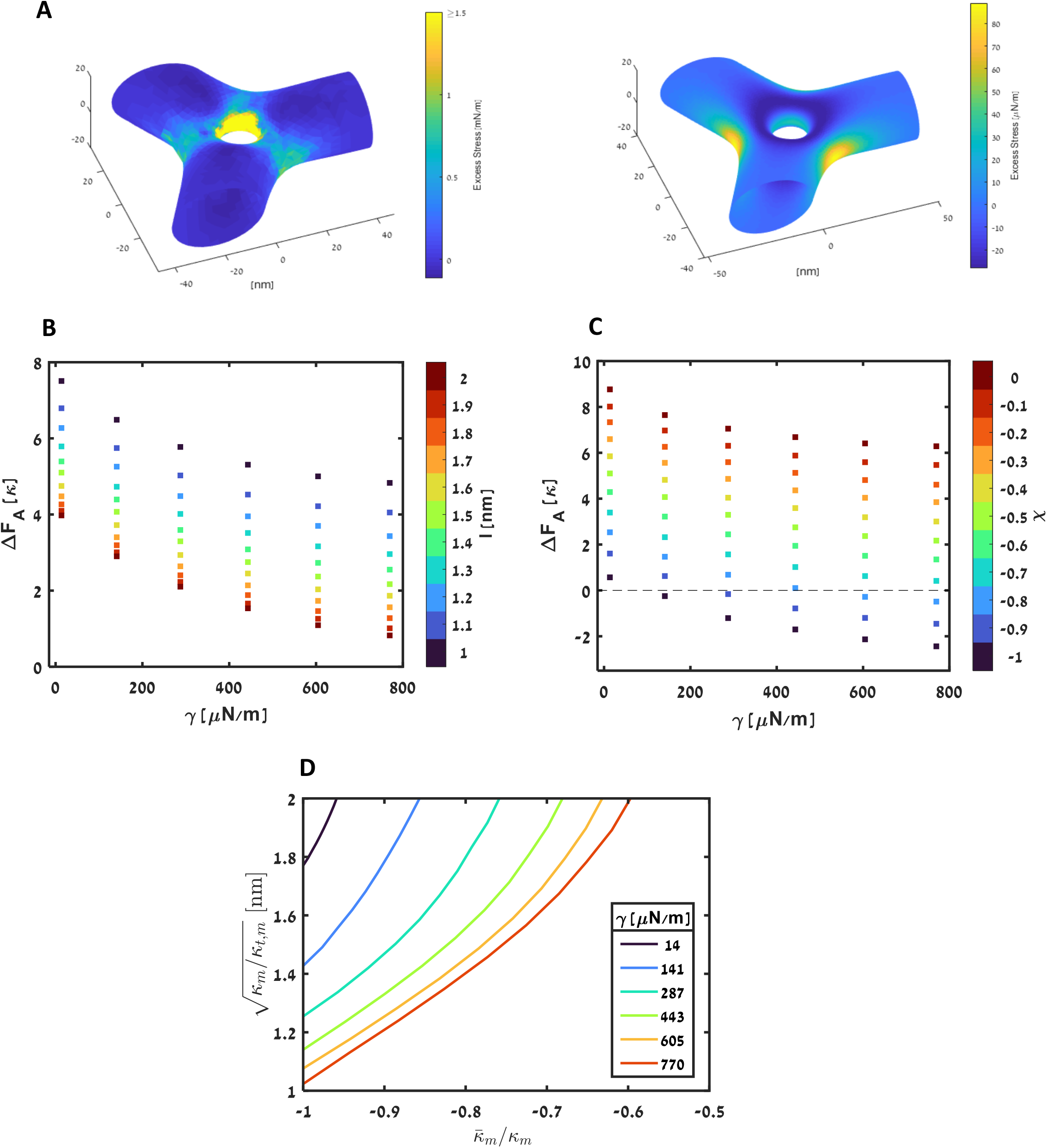
Hemi-fission stage of the pore sealing in a junctional knot. (A) The excess stress distribution along the membrane of a junctional knot for two characteristic values of the ratio between the membrane modulus of Gaussian curvature and bending modulus, 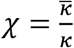, and other elastic parameters as in (Fig.2). Left panel, *χ* = −1; right panel *χ* = 0. (B) The activation energy of hemi-fission as a function of the membrane tension, *γ*, for 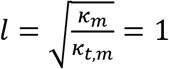, and different values of 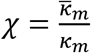. (C) The activation energy of hemi-fission as a function of the membrane tension, *γ*, for *χ* = −0.5 and different values of *l*. (D) The relationships between 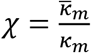 and 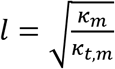 corresponding to a vanishing activation energy of hemi-fission, Δ*F_A_* = 0, for different values of the tension.

Based on the previous work(Kozlovsky and Kozlov, 2003), we consider the fission reaction to proceed via an intermediate stage of hemi-fission at which only the internal monolayer of the pore rim undergoes splitting, while the external monolayer remains intact, as illustrated in (Fig. 1E-F). The intermediate structure formed at this stage is referred to as the hemi-fission stalk(Kozlovsky and Kozlov, 2003) (Figs. 1F). The fission is completed by splitting of the hemi-fission stalk (Figs. 1G,H). Since, as shown above, the full fission results in a decrease of the system elastic energy, we consider the hemi-fission stalk to represent the potential rate-limiting step of the fission process. The activation energy of fission, Δ*F_A_*, is thus the difference between the energy of a junctional knot with the hemi-fission stalk, *F_EHF_*, and that of the initial junctional knot with a pore, *F*_*E*1_, such that Δ*F_A_* = *F_EHF_* – *F*_*E*1_.

The computation of the configuration and the energy of the hemi-fission stalk is based on the previous works (Hamm and Kozlov, 2000) and, as already mentioned, uses the elastic model of tilt, splay and saddle-splay deformations of lipid monolayers described in (SI B). The monolayer resistance to the three deformations is characterized by the elastic moduli of tilt, *κ_t_*, splay, *κ_m_*, and saddle-splay, 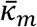 (SI B). As described in (SI C), the computational procedure we used here is similar to that developed previously for modeling intermediate structures of different reactions of membrane fusion and fission(Golani et al., 2021; Kozlovsky et al., 2002; Kozlovsky and Kozlov, 2002), but accounts for the specific geometry of a junction between three tubules.

Our main goal was to examine the dependence of the activation energy, Δ*F_A_*, on the membrane tension, *γ*, which we took to vary in the range between 14 and 770 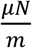. We also explored the dependence of the activation energy, Δ*F_A_*, on the monolayer elastic parameters whose values have not been directly measured and can vary within estimated ranges. Specifically, we computed Δ*F_A_* for different values of the dimensionless ratio between the monolayer modulus of Gaussian curvature and the bending modulus, 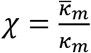, varying in the feasible range between 0 and −1(Templer et al., 1998). Another parameter is the characteristic decay length of the tilt deformation, *l*, which is determined by the ratio between the monolayer bending and tilt moduli by, 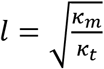, and was taken to change between 1 and 2nm(Golani et al., 2021).

The results for the energy of the hemi-fission intermediate, *F_EHF_*, are shown in (Fig. 3A), and for the activation energy, Δ*F_A_*, are presented in (Fig. 3B, 4B-C), where the energy values are scaled by the bilayer bending modulus *κ* ≈ 10^−19^Joule(Dimova, 2014), the reference system is the same as used for computations of the junctional knot energy (Fig. 1J), and the tension varies within a biologically feasible range.

The dependence of Δ*F_A_* on the tilt decay length, *l*, is shown in (Fig. 4B) for *χ* = −0.5. The activation energy is decreasing with increasing, *l*, which means that the smaller the tilt modulus, *κ_T_*, the smaller the activation energy of fission.

The dependence of Δ*F_A_* on *χ* is presented in (Fig. 4C) for *l* = 1.5 nm. The activation energy decreases with *χ*, and, consequently, with the monolayer modulus of Gaussian curvature 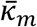, becoming more negative. For sufficiently negative values of *χ*, the activation energy is predicted to be small or vanishing.

The effects of the elastic parameters and tension on the activation energy are summarized in (Fig. 4D), where each line represents the parameter combinations for which the activation energy vanishes meaning that the fission process happens spontaneously. According to this diagram, for fission to be spontaneous, the tension has to be large, the monolayer modulus of Gaussian curvature has to be as negative as possible, and the tilt modulus has to be small.

The computed membrane configurations formed along the full pathway of the junctional knot remodeling into a regular junction are presented in (Fig. 5A-C).

**Figure 5.**
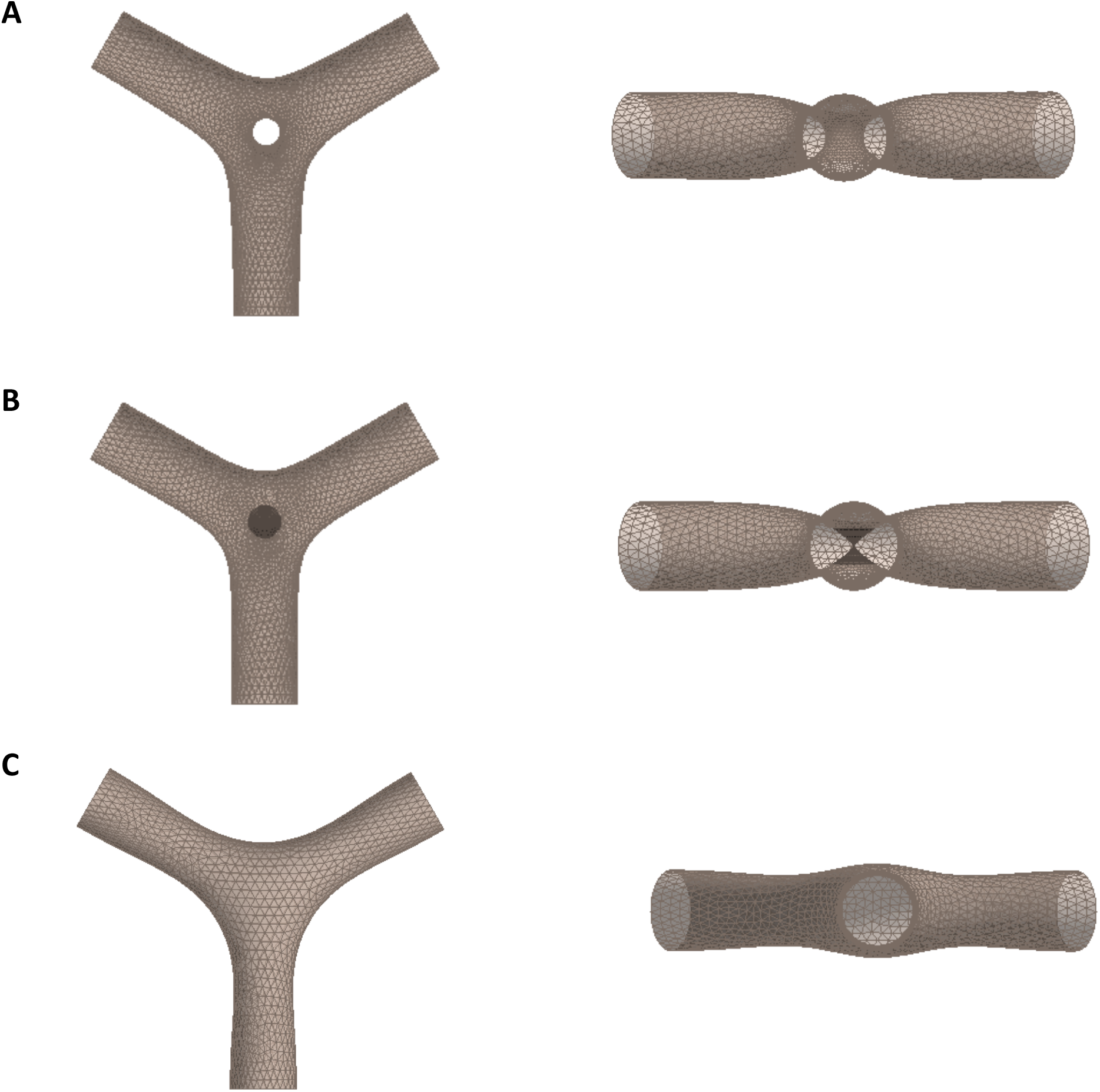
Computed configurations of all stages of a junctional knot transition into a regular junction with the top and plain views presented, respectively, in the left and right panels. (A) Junctional knot with one pore. (B) Hemi-fission stage of the pore sealing. (C) Regular junction. *κ* = 0.8 · 10^−19^ *J*; 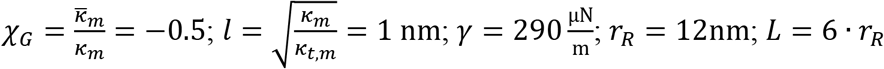.

## Discussion

Here we considered the critical final stages of the ER ring closure, the process of constriction and vanishing of polygonal unit-cells of the ER tubular networks. The ring closure reduces the number of the network three-way junctions, which dynamically equilibrates the persistent process of new junction generation by the ER tubule branching and fusion. We computationally analyzed the experimentally inaccessible structures of the junctional knots, which result from constriction of one or several triangular unit-cells of a tubular network and contain internal pores. We analyzed the pore sealing by membrane fission, which completes the ring closure, and determined the conditions necessary for vanishing of the activation energy related to the hemi-fission intermediate of the fission reaction.

The major message of our analysis is that the ER ring closure can be driven by tension existing in the ER tubular membranes. The distribution of the membrane tension in ER appears to be highly variable. According to the ER images in live cells, there are regions where the ER tubules look loose and undergoing thermal undulations(Georgiades et al., 2017; Nixon-Abell et al., 2016; Westrate et al., 2015), which implies that they are subjected to either vanishing or extremely low tensions(Georgiades et al., 2017). At the same time, the ER tubules, which undergo sliding and ring closure look stretched(Lee and Chen, 1988; Powers et al., 2017), as expected for membranes exposed to considerable tensions. Substantial tensions in reconstituted ER networks were determined by direct measurements(Upadhyaya and Sheetz, 2004). Altogether, these observations suggest that different parts of the ER network are subject to various tensions, which is counterintuitive given the 2D-fluidity of the interconnected ER membranes. Yet such situations are feasible provided that not the whole ER tubular network but only certain parts of it are stretched between the sites of contact with cytoskeleton in which pulling forces are applied to the ER tubules. While these ER areas must be exposed to tension, the other ER regions, which are outside of the zones of the pulling force action, are expected to remain tension free.

According to our results, the spontaneous pore sealing corresponding to a vanishing activation energy of the membrane fission reaction can be driven by small membrane tensions generated by pulling forces as little as *f* = 2 *pN* if the absolute value of the membrane modulus of Gaussian curvature is large, e.g. 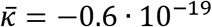 *Joule*, while the tilt modulus is small such as 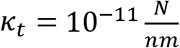. Such parameters values correspond to the limits of the currently estimated ranges for 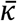 and *κ_t_*. For the mid-range values of, 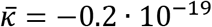 *Joule* and 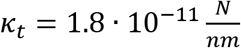, the spontaneous fission requires large tensions corresponding to pulling forces of about *f* = 40 *pN*. Such tensions could be generated by more than 10 molecular motors acting on each membrane tubule, which must be, biologically, a seldom situation. This implies that special fission proteins might be required to facilitate the junctional knot conversion into the regular junctions.

Another prediction of our results is that in ER network regions exposed to ultralow tensions, the transition of the junctional knots to regular junctions should not occur. In this case, one might expect the accumulation of pores within the junctional knots leading to formation of sieve-likes structures such as ER matrices(Nixon-Abell et al., 2016).

## Acknowledgements

The authors are grateful to Tom Rapoport for stimulating discussions. MMK was supported by Deutsche Forschungsgemeinschaft (DFG) through SFB 958 “Scaffolding of Membranes”, and Israel Science Foundation grant 3292/19, and holds Joseph Klafter Chair in Biophysics.

## Supplementary Information

### A. Relationship between the radius and tension of a reservoir tubule

To derive the relationship between the radius, *r_R_*, the tension, *γ*, and the spontaneous curvature, *J_S_*, of a tubular membrane given by (Eq. 2) of the main text, we use the equation of mechanical equilibrium of a curved membrane (see “Some aspect of membrane elasticity”(Poon and Andelman, 2006, pp. 79–96)) Since the mean curvature, *J*, of a cylindrical membrane is homogeneous so that its gradient along the membrane surface vanishes, ∇*J* = 0, and the Gaussian curvature vanishes as well, *K* = 0, the equilibrium equation adopts a simple form,

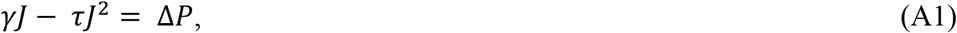

where *τ* is the membrane torque related to the membrane spontaneous curvature, *J_S_*, and bending modulus, *κ*, by the relationship

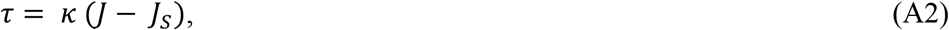

and Δ*P* is the trans-membrane pressure difference. Substituting (Eq.A2) into (Eq.A1) and taking into account that, Δ*P* = 0, for ER tubules, we obtain the equation for the tubule mean curvature

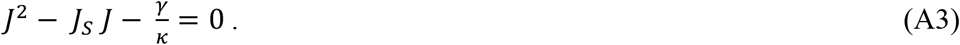

Solution of this equation for the tubule mean curvature, *J*, and cross-sectional radius, *r*, gives

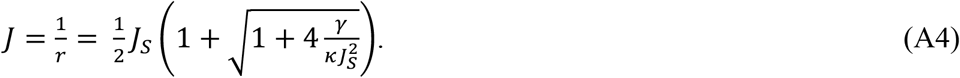

For vanishing spontaneous curvature, *J_S_* = 0, the expression (Eq.2) takes a simple form

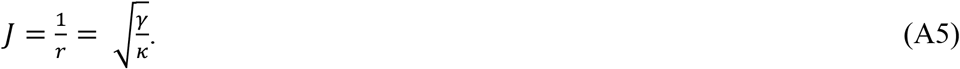

The relationships (Eqs. A4,A5) describe the curvature, *J_R_*, and the cross-section radius, *r_R_*, of the reservoir given by (Eq. 2) of the main text.

### B. Elastic energy of splay and tilt of membrane monolayer

Hemi-fission and hemi-fusion intermediates of membrane remodeling introduce deformations of inhomogeneous tilting of the hydrocarbon chains of lipid molecules(Hamm and Kozlov, 2000, 1998). Since the in-plane distributions of these deformation are different for the two monolayers, the elastic energy has to be computed separately for each monolayer, which can be done by using the generalized Helfrich model accounting for the tilt, splay, and saddle splay of the hydrocarbon chains as introduced in detail in the earlier works(Kozlovsky et al., 2004, 2002; Kozlovsky and Kozlov, 2003, 2002). Here we sketch the main features of this model.

To describe the monolayer geometry, we use the dividing surface coinciding with the interface between the regions of the polar heads and hydrocarbon tails of the monolayer. The local orientation of the lipid molecules at each point of the dividing surface is characterized by the lipid director, 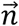, which is defined as a unit vector pointing from the locally averaged position of the ends of the hydrophobic lipid tails to those of the polar heads (Fig. S1). Lipid tilt, splay, and saddle splay are defined based on the variation of 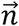 along the dividing surface and the deviation of 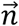 from the unit normal vector to the dividing surface, 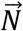.

The lipid tilt is defined as

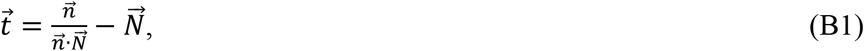

and its consequence for the monolayer internal structure can be understood as shearing of the lipid hydrocarbon chains in the direction perpendicular to the monolayer plane(Hamm and Kozlov, 2000, 1998; May et al., 2004) (Fig. S1).

Lipid splay, 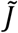, referred also to as the modified mean curvature, is defined as the two-dimensional divergence of the lipid director, 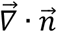, determined along the, generally, curved dividing surface. In covariant form, the lipid splay is the trace of the tensor, 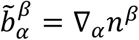, where the sub and superscripts denote, respectively, the co- and contravariant components in the local coordinate basis of the surface. 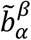 is referred to as the modified curvature tensor. Therefore,

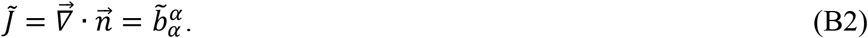

Lipid saddle splay is the determinant of the modified curvature tensor,

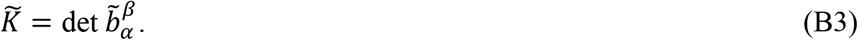

In the absence of the tilt deformation, the lipid director and the surface normal are aligned, 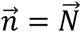, so that the splay, 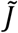, is reduced to the two-dimensional divergence of the surface normal, 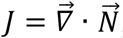, whose geometrical meaning is the mean curvature of the dividing surface, *J* = *c*_1_ + *c*_2_, where *c*_1_ and *c*_2_ are the two principal curvatures of the surface. Saddle splay, 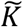, is reduced in this case to Gaussian curvature of the surface, *K* = *c*_1_ · *c*_2_, the product of the two principle curvatures.

The elastic energy of the tilt, splay and saddle-splay deformations per unit area of the membrane plane is given by(Hamm and Kozlov, 2000),

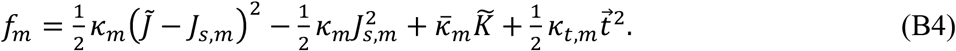

where *κ_m_* is the monolayer bending modulus whose value was measured for pure lipid monolayers and varies in the range between 5-20 *k_b_T* (Dimova, 2014; Niggemann et al., 1995; Pontes et al., 2013) (*k_b_T* ≈ 4.11 · 10^−21^ *Joule* is the product of the Boltzmann constant, *k_B_*, and the absolute room temperature); 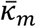 is the monolayer saddle splay modulus, which has been estimated to be negative and have an absolute value smaller but of the same order of magnitude as the bending modulus; *κ_t,m_* is the monolayer tilt modulus; *J_s,m_* is the monolayer spontaneous splay, which equals the molecular intrinsic curvature of the constituting lipids. We define the ratio between the bending and saddle splay moduli as 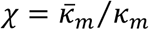, which can have values in the range between −1 and 0(Templer et al., 1998; Terzi et al., 2019). As a measure of the tilt modulus, we use 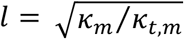, which has a meaning of a characteristic decay length of the tilt deformation and whose estimated values vary in the range between 1nm and 2nm(Doktorova et al., 2017; Hamm and Kozlov, 1998; May et al., 2004; Nagle et al., 2015; Terzi et al., 2019). The ER membrane is composed mostly of phosphatidylcholine (PC) and phosphatidylethanolamine (PE)(Zambrano et al., 1975), both having negative intrinsic molecular curvature(Chen and Rand, 1997; Leikin et al., 1996; Szule et al., 2002), and a small amount of lysolipids, such as lysophosphatidylcholine (LPC), characterized by positive intrinsic molecular curvature (Fuller and Rand, 2001; Zambrano et al., 1975). We estimate the ER monolayers spontaneous splay by calculating the weighted average of the intrinsic molecular curvatures of the constituting lipids to be *J_s,m_* = −0.1 *nm*^−1^ (Kozlov and Helfrich, 1992). Since, to the best of our knowledge, there is no evidence for asymmetrical distribution of lipids components between cytosolic and luminal monolayers of ER membrane, we assume the two monolayer compositions to be identical.

The total elastic energy of the membrane is calculated by integrating the energy density (Eq. 4) over the areas of the luminal and cytosolic monolayers,

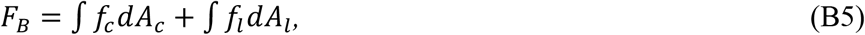

where the subscripts *c* and *l* denote the cytosolic and luminal monolayers of the ER tubule, respectively. The energy of membrane tension, *F_T_*, is given by a product between the lateral tension of each membrane monolayer, *γ_m_*, and its area, which is *A_c_* and *A_l_* for the cytosolic and luminal area respectively,

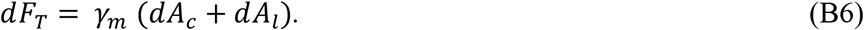

The total elastic energy is:

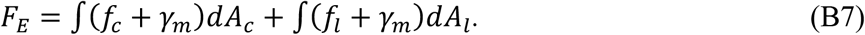

### C. Structure and energy of the hemi-fission stage of the pore sealing

We consider the pore sealing to proceed through an intermediate stage of hemi-fission of the membrane neck constituting the rim of the pore (Fig.1E-H of the main text). The hemi-fission intermediate includes a non-bilayer structure in the middle (Fig. 1F of the main text) referred to as the membrane stalk. Our goal here was to compute the elastic energy of the hemi-fission intermediate, *F_EHF_*, which could be compared to the energies of a junctional knot with one pore, *F*_*E*1_, and of a regular junction with no pore, *F*_*E*0_, as described in the main text.

To perform the computation of *F_EHF_*, we used the results of the previous works(Kozlovsky and Kozlov, 2003, 2002) in which the structure and energy of a membrane stalk were computed for various processes of membrane remodeling: abscission of an endocytic vesicle(Kozlovsky and Kozlov, 2003), hemi-fusion of flat(Kozlovsky et al., 2002; Kozlovsky and Kozlov, 2002) and curved(Martens et al., 2007) membranes. In spite of different overall configurations of these systems, the computed structure of the membrane stalk had a common feature: the middle part of the stalk could be considered as a localized sub-system referred to as the stalk-core (Fig.S1). The tilt angles in the stalk-core monolayers were equal 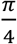 in the middle of the structure, and relaxed along the mid-plane over a characteristic length of few nanometer(Kozlovsky and Kozlov, 2003), which sets the stalk-core boundary (Fig.S1).

Based on that, we approximated the hemi-fission intermediate of the pore sealing as the stalk-core embedded into the three-way tubular junction. The boundary between the stalk-core and the surrounding membrane was chosen to lie on a sphere co-centered with the stalk-core, having a radius as *R_b_* = 8 *nm*, and called below as the boundary sphere. According to the earlier works(Kozlovsky and Kozlov, 2003, 2002) and verified by the present computations for the stalk-core monolayers, at this boundary the lipid tilt, practically, vanishes and the monolayer curvatures becomes small with respect to the monolayer width.

The computation of the configuration and energy of the hemi-fission intermediate included the calculation of the stalk-core energy and the energy of the surrounding membrane of the junction and minimization of the sum of the two energies with respect to the position of the interface between the stalk-core and the surrounding membrane on the boundary sphere.

By computing the stalk-core energy, we assumed that reticulons are excluded from the stalk-core because of the small dimension and non-bilayer structure of the latter so that the spontaneous splay of the stalk-core monolayers is determined solely by its constituting lipids. We further assumed that the monolayer hydrocarbon moiety is incompressible and that the monolayers dividing planes are parallel to the membrane mid-plane, meaning the lipid tail length is related to the tilt by the relationship 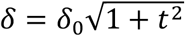. The computation of the stalk-core energy consisted in integration of the monolayer energy density along the monolayer surfaces (Eq. B7) and minimization of the obtained overall energy with respect to the tilt distribution in each monolayer upon a requirement of the monolayer tilt angles remaining equal 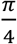 in the center of the structure (Fig.S1). The details on the parametrization of the monolayer shapes and tilt distributions we used here can be found in(Golani et al., 2021).

The computation of the energy of the surrounding membrane was performed analogously to that of the junctional knot or a regular junction, i.e. by integration of the regular density of the bilayer elastic energy ((Eq.3) of the main part) and minimization of the result with respect to the membrane shape using the Brakke’s Surface Evolver(Brakke, 1992).

The computations of the stalk-core and the surrounding membrane energies were coupled through fulfillment of the boundary conditions at the interface between these two parts of the system, which included the requirement of smoothness of the membrane mid-plane profile, the continuity of the mid-plane curvature and the vanishing of the lipid tilt in the two monolayers of the stalk-core.

## Figure legend

**Figure S1.**
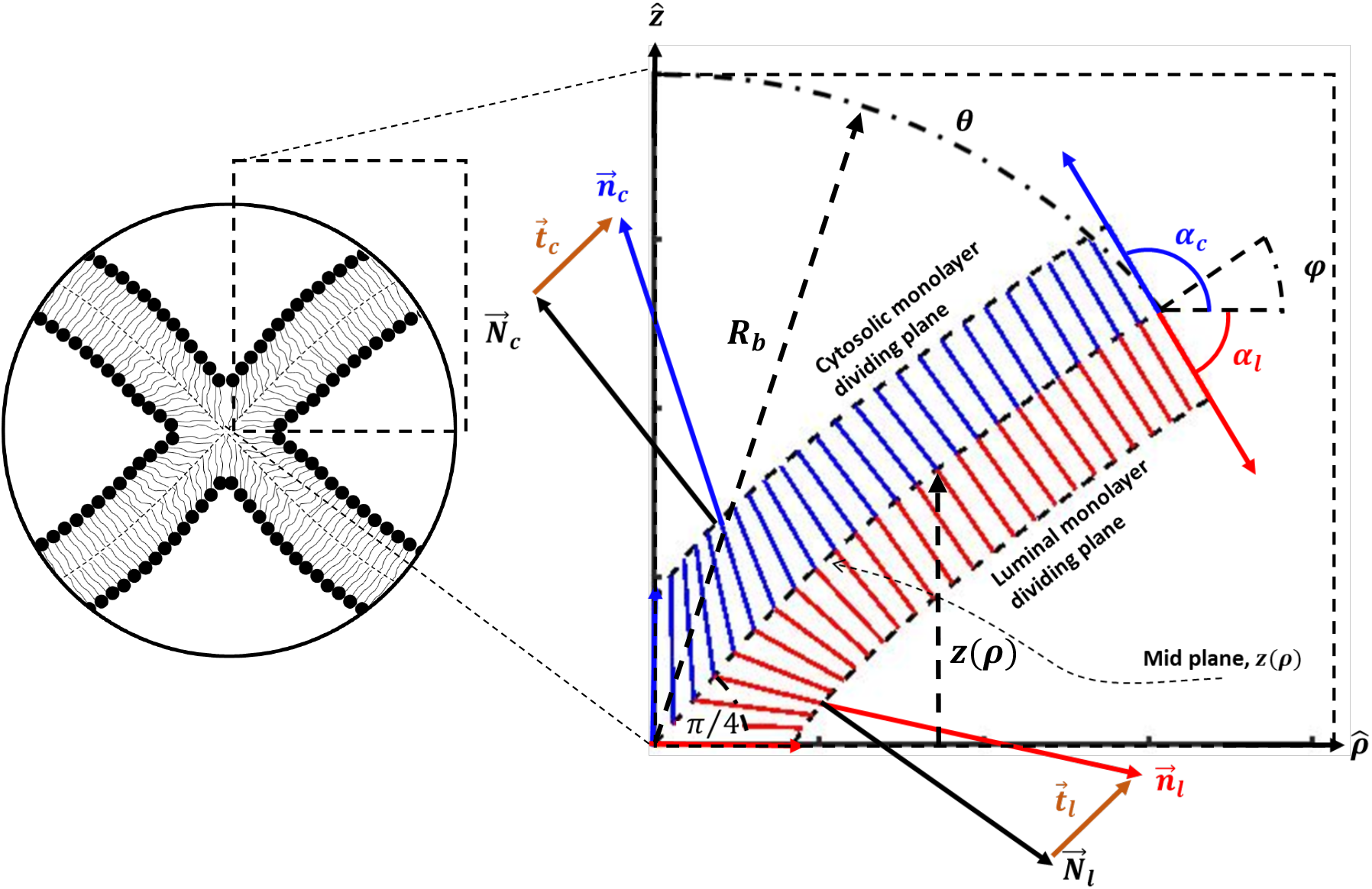
Hemifission stalk and definitions. The vertical, *Z*, axis represents the axis of rotational symmetry. Left – drawing of a cross section of a hemifission stalk, bordered with by a sphere representing the interface e between the stalk-core and the surrounding membrane. right – computation results of stalk core shape. Blue and red lines represent averaged lipid director of cytosolic and luminal monolayers, respectively. Definitions: 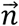 lipid director, 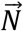 Normal to dividing plane, 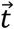 tilt vector, *α* angle lipid director and 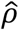 and *φ* angle between tangent vector to mid-plane and 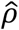. The subscript *c* and *l* represent the cytosolic and luminal monolayers, respectively. Right panel represent the minimal energy configuration for the parameters: = 1.5*nm, χ* = −0.5, *J_m_* = −0.0*nm*^−1^, *θ_b_* = 50°, *φ_b_* = 31° and *R_b_* = 8*nm*.

